# Inference of cell division parameters in bacteria under transient growth conditions

**DOI:** 10.64898/2026.11.05.764660

**Authors:** César Nieto, Sayeh Rezaee, Cesar Augusto Vargas-García, Abhyudai Singh

## Abstract

Most quantitative models of bacterial cell division assume steady-state growth, yet natural environments are highly dynamic. Here, we infer division control parameters of *E. coli* from single-cell size trajectories under nonsteady growth. We describe growth as a continuous process interrupted by stochastic divisions, whose propensity depends on cell size and on the size added since birth. Combining an estimation of time-dependent single-cell growth rates with a maximum-likelihood estimator that we previously validated on synthetic data, we obtain time-resolved estimates of the mean added size at division, its noise, and a strategy parameter that measures how added size depends on birth size. This parameter vanishes for the *adder*, in which cells add, on average, a fixed size regardless of their size at birth, and is negative for *sizer-adder* control, in which larger-born cells add less size. We apply this approach to published single-cell data covering steady exponential growth, gradual nutrient depletion, and exit from starvation. We implement a time-resolved model selection based on the Akaike information criterion, concluding that the adder is sufficient during steady growth, whereas size dynamics during entry into and exit from stationary phase are better explained by transient deviations toward sizer-adder control. The framework provides a general method to quantify time-resolved changes in the scale, stochasticity, and strategy of division in nonsteady environments.

## I. INTRODUCTION

Cell size regulation is a fundamental process through which cells maintain physiological function and adapt to diverse growth conditions [1–4]. To keep their size within a characteristic range, proliferating cells must balance how fast they grow with how often they divide [5– 9]. Growth itself integrates many internal processes, including nutrient sensing [10, 11], the regulation of ribosome abundance [12], the rates of cell elongation and DNA replication [13, 14], and physiological responses to environmental factors such as pH [15]. The processes that determine cell division are tightly interconnected and remain an active area of study [16, 17]. Quantifying division is important because division sets not only cell size regulation but also the proliferation rate and, hence, how successfully a population adapts to its environment [18–21].

Most quantitative characterizations of division control have been obtained under steady growth, where size statistics are stationary and comparatively easy to analyze [22]. Natural habitats rarely provide such constancy: nutrients are consumed, and pH, osmolarity, or antibiotic concentrations change over time [23– 25]. Under these conditions, size regulation must be readjusted continuously. Such responses can now be followed directly by combining microfluidic trapping with long-term single-cell imaging [26]. These measurements show that bacterial size changes strongly with the environment: cells are large in rich media and shrink as nutrients become limiting [27, 28]. One proposed explanation is that, when resources fluctuate, size reflects how cells partition their proteome between biomass synthesis and the division machinery [29, 30]. Linking these size changes to a specific division mechanism is nonetheless difficult, because the molecular players and their regulation are not conserved across species [31, 32]. In this situation, *coarse-grained* models that summarize division control with a few effective parameters provide a common language to compare organisms and conditions without committing to a particular molecular mechanism [31, 33].

At this coarse-grained level, size control is usually described with reference to three division strategies [31, 34–38]. In a *sizer*, division is triggered once the cell reaches a threshold size; in an *adder*, the size accumulated between birth and division is, on average, the same regardless of birth size; and in a *timer*, the cell cycle lasts a fixed time irrespective of size. The three are conveniently distinguished by regressing the added size at division on the size at birth. The slope of this regression equals −1 for a sizer, 0 for an adder, and +1 for a timer. Intermediate strategies are called *sizer-adder* (slopes in (−1, 0)) and *timer-adder* (slopes in (0, 1)) [39–41].

Because this regression is built from completed cycles, it requires pooling many birth–division pairs, typically collected over long periods of balanced growth [42]. When the environment changes within a generation, pooling mixes cycles that experienced different conditions, and there is no natural way to assign a strategy to a given instant. An alternative is to describe division through a propensity, or hazard rate: the probability per unit time that a cell divides given its current state [43–45]. The propensity can be interpreted as an effective summary of hidden cell-cycle progression, for instance the build-up of division regulators such as FtsZ, and it is defined at every instant [19]. Model parameters can therefore be allowed to vary in time together with the environment.

In previous work, we introduced a family of division propensities that depend on cell size and on the size added since birth, and showed how their parameters map onto the classical division strategies [33]. We then derived a maximum-likelihood estimator for these parameters, including a strategy parameter that interpolates between sizer-adder and timer-adder control, and benchmarked it on synthetic trajectories with prescribed time-varying parameters [46]. Here we apply this framework to experimental data under nonsteady growth. Using published microscopy tracking measurements of *E. coli* [27], we analyze three regimes: balanced growth in rich medium, nutrient depletion, and exit from starvation after fresh medium is supplied. Beyond the application itself, we (i) incorporate time-dependent single-cell growth rates estimated nonparametrically from the size trajectories, (ii) combine the estimator with time-resolved model selection based on the Akaike information criterion, which tests in each time window whether departures from the adder are statistically supported, and (iii) compare the continuous-time inference with the conventional per-cycle regression. We find that division remains close to an adder during balanced growth but shifts transiently toward sizer-adder control as when cells recover from slowdowns and during the first divisions after starvation, while the target added size follows the successive slowdowns of growth.

The paper is organized as follows. Section II summarizes the stochastic model of growth and division, the inference procedure, the data set, and the estimation of growth rates. Section III presents the per-cycle and time-resolved analyses, and Sec. IV discusses their physiological interpretation and the limitations of the approach.

## II. MODEL AND INFERENCE FRAMEWORK

We first summarize the mathematical basis of the inference method; further details are given in Ref. [46].

### A. Cell size dynamics

We use cell length, denoted by *s*(*t*), as a measure of cell size. The added size, ∆(*t*) = *s*(*t*) − *s*_*b*_, is the increase in size since the beginning of the current cycle, where *s*_*b*_ is the size at birth; hence ∆ = 0 at birth and grows throughout the cycle. Between divisions both variables evolve as

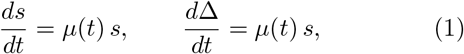

where *µ*(*t*) *>* 0 is the single-cell growth rate, which may vary in time and is estimated from the size trajectories (Sec. II H).

### B. Division propensity

Divisions are treated as discrete stochastic events superimposed on continuous growth: the probability that a cell divides during an infinitesimal time interval (*t, t* + *dt*] is *h*(*µ, s*, ∆) *dt*, where *h* is the division propensity. An adder is obtained when the propensity factorizes as [33, 46]

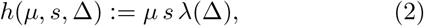

where *λ*(∆) *>* 0 is the *division function*. For the adder, it depends on the added size alone; other strategies follow when *λ* also depends on *s*.

Integrating the propensity over a finite window *δt* (typically the interval between consecutive frames, short compared with the cell cycle but not necessarily infinitesimal) gives the probability of observing a division between consecutive measurements. For a cell with size *s*(*t*_0_) and added size ∆(*t*_0_) at time *t*_0_,

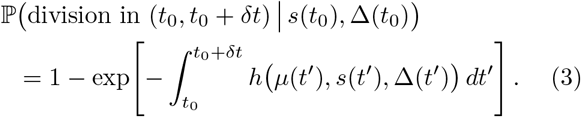

This probability forms the basis of the maximum-likelihood inference described below. We first introduce analytical approximations that simplify the evaluation of the integral.

### C. Sigmoidal division function

Because the molecular state of the division machinery is not observed, the functional form of *λ* is not fully determined *a priori* ; it can be derived from a specific mechanism [33] or chosen phenomenologically [47]. We use a sigmoidal division function, an analytically tractable form whose stochastic properties closely match those of more complex alternatives [48]:

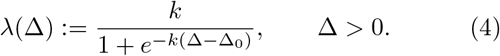

Here ∆_0_ is the scale parameter and *k* the shape parameter. With this division function, one can compute the moments of the added size at division ⟨∆d⟩, namely its mean ⟨∆d⟩ and its noise, quantified by the squared coefficient of variation 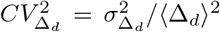 , where 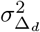 is the variance of ⟨∆d⟩ [33]. For weakly noisy division 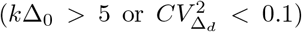 the two parameters relate to the measurable moments as [46, 49]

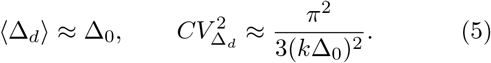

If growth within each window is approximated as exponential at a constant rate *µ* (which may change from one window to the next), the integral in Eq. (3) can be evaluated in closed form [46]:

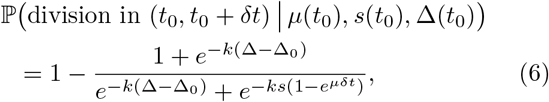

where the parameter vector *θ* := (∆_0_, *k*) is related to the division statistics ⟨∆d⟩ and 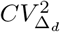 through Eq. (5).

### E. Maximum-likelihood estimation

At each acquisition time *t*_*n*_, the data consist of the triplets (*µ*_*i*_, *s*_*i*_, ∆_*i*_), that is, the growth rate, size, and added size of the *i*th cell at *t*_*n*_. Depending on whether the cell divides before *t*_*n*+1_, it contributes a log-likelihood

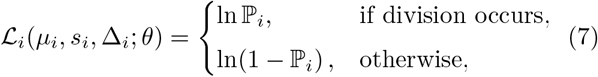

where P_*i*_ is the division probability of the *i*th cell given by Eq. (6), evaluated at the measured values (*µ*_*i*_, *s*_*i*_, ∆_*i*_). The parameters are inferred by maximizing the summed log-likelihood,

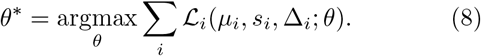

Each term involves only the instantaneous values of *s*, ∆, and *µ*, with no explicit reference to the birth size or to earlier times. The description is therefore Markovian in the state (*µ, s*, ∆) and fully characterize by the time-dependent division parameters *θ*^*∗*^ = (∆_0_, *k*) or, equivalently, 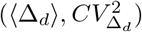 The estimate can thus be computed independently in every time window, under the single assumption that cells observed within the same window share the same division parameters.

### E. Alternative division strategies

The approach extends to strategies beyond the adder (timer-adder and sizer-adder). To this end, we take the mean added size at division to depend linearly on birth size *s*_*b*_ [46, 50–52]:

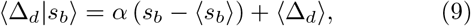

where ⟨∆d⟩ *s*_*b*_ is the mean added size conditioned on birth size and *α* is the strategy parameter: *α* = 0 for the adder, −1 *< α <* 0 for sizer-adder, and 0 *< α <* 1 for timer-adder control. The noise 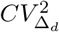 could also depend on *s*_*b*_, but for simplicity and numerical stability we take it to be constant [46]. The division probability is then computed from Eq. (6), with parameters approximately given by

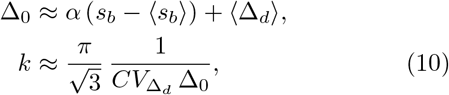

which follows from Eq. (5) and holds for weakly noisy division (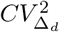*<* 0.1). Using *s*_*b*_ = *s* − ∆ and the approximation *s*_*b*_ ⟨∆d⟩ , the propensity depends explicitly only on *s*, ∆, and *µ*, and parametrically on the interpretable statistics 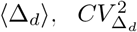, and *α*. Equation (10) is shown for illustration only; the full dependence is more complex and is detailed in Ref. [46].

### F. Model comparison

In each time window, we fit both the adder model (*α* = 0, with two parameters: ⟨∆d⟩ and 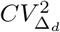) and the alternative model (*α* 0, with three parameters: ⟨∆d⟩, 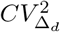, and *α*). The two models are compared using the Akaike information criterion (AIC) [53],

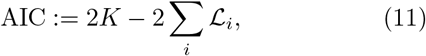

where *K* is the number of fitted parameters and the sum is evaluated at the maximum-likelihood estimate *θ*^*∗*^. We denote the criterion by AIC_*α*=0_ for the adder and by 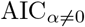 for the alternative model.

With AIC_min_ = min(AIC_*α*=0_, AIC_*α*=0_), the relative likelihood of the adder model is

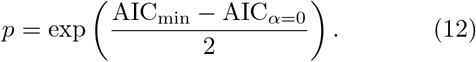

Values of *p* close to 1 indicate that the simpler adder model is supported by the data, whereas small values indicate weaker support for the adder. Based on *p*, our approach identifies the time windows in which the adder model suffices and those in which a deviation (*α* 0) is statistically favored. We consider the alternative model favored when *p <* 10^−3^.

### G. Experimental data

Single-cell growth and division data were obtained from Ref. [27], in which *E. coli* MG1655 lineages were tracked in a *mother machine* [Fig. 1(a)], a microfluidic device that traps a mother cell at the closed end of each channel while its descendants are washed away [22].

**FIG 1.**
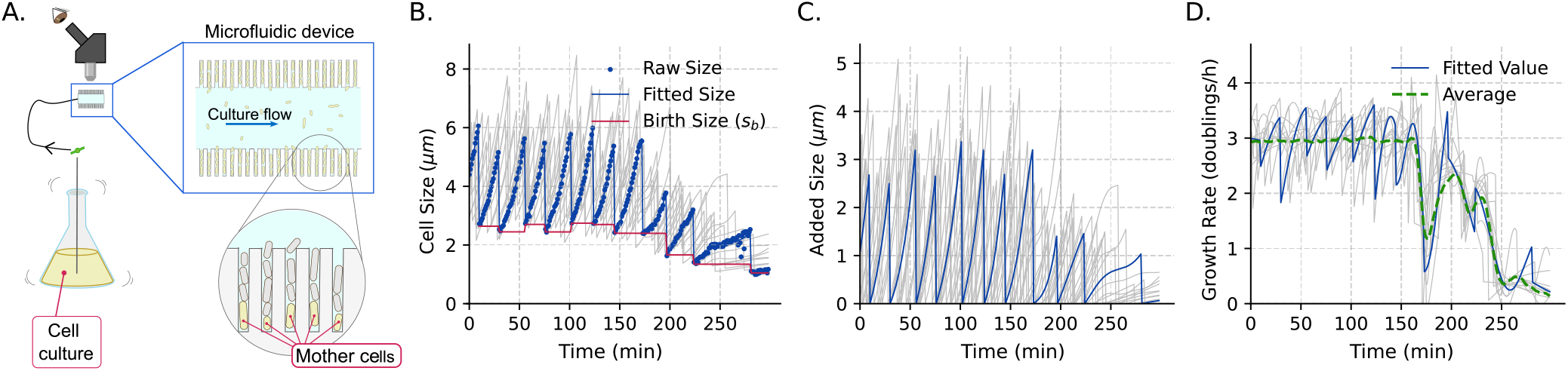
Experimental setup and variables measured for single-cell size tracking. (a) Schematic of the experimental setup used to generate the published data set [27] analyzed in this study. *E. coli* cells (mother cells) were trapped at the closed ends of the channels of a mother-machine microfluidic device and subjected to dynamically controlled nutrient flows. During steady growth, fresh medium was continuously supplied to the device. During entry into stationary phase, the channels were fed from a batch culture in a flask; as nutrients were depleted in the batch culture, the trapped cells underwent a metabolic downshift. To capture the exit from stationary phase, previously starved cells were supplied with a sudden influx of fresh medium to resume growth. (b) Cell length trajectories over time. A representative single-cell trajectory is highlighted, showing raw length measurements (blue dots), fitted cell length (solid blue line), and estimated birth size *s*_*b*_ for each division cycle (red line). Gray lines show the length trajectories of other individual cells. Data and growth rate (c) Added size dynamics, defined as the increase in cell length relative to birth length within each cell cycle. The representative cell (blue line) is shown together with the added size of other cells (gray). (d) Instantaneous growth rates of individual cells (gray) and the representative single cell (blue line), shown alongside the mean growth rate across tracked cells (green dashed line), measured in doublings per hour. Growth rates were estimated using Gaussian process regression with a time-varying growth rate.

Cells were grown at 37 ^*◦*^C in EZ-rich defined medium (EZRDM), with the device coupled to shaking batch cultures so that trapped cells experienced the same time-varying nutrient conditions as the bulk population. We analyzed three conditions: (1) steady exponential growth in fresh EZRDM at constant growth rate; (2) entry into stationary phase, in which cells experience progressive nutrient depletion, passing through diauxic shifts before growth arrest; and (3) exit from stationary phase, in which growth resumes after fresh medium is introduced. Phase-contrast images acquired at 1-min intervals captured 952 individual lineages over more than 30 h. Image segmentation and cell tracking produced cell-length trajectories, which were manually verified to refine division times. From these trajectories we extracted the birth length and the added length (division length minus birth length) of each complete cycle [Figs. 1(b) and 1(c)].

### H. Growth rate inference

After verifying manually the division time events, it is possible to fit the cell size dynamics between divisions to smooth functions of time. Instantaneous single-cell growth rates were estimated with a nonparametric method based on Gaussian processes (GPs) [54]. This approach assumes the growth rate to be locally exponential changing smoothly over time. If *s*(*t*) is modeled as a GP, its first and second time derivatives are also GPs, so that smoothed estimates of *s* and *ds/dt* can be obtained directly from the time series. Throughout, *µ* = *d* ln *s/dt* is the exponential growth rate (in min^−1^ in the equations); figures report it in doublings per hour, 60 *µ/* ln 2. After fitted *s*, ∆ and *µ*, we use these estimates to implement the MLE of division parameters.

## III. RESULTS

In the first part of the analysis, we applied the classical per-cycle approach based on the original study [27], in which cycles are indexed not in continuous time but by their order relative to entry into or exit from stationary phase. We then used the occurrence of divisions within each 1-min window to infer the propensity parameters by maximum likelihood, and compared the adder with alternative strategies using the AIC. This analysis provides a more detailed characterization of size regulation under time-varying growth rates.

### A. Classical cell-cycle analysis of division strategies during nutrient shifts

In the classical discrete-event cell-cycle analysis, complete cell cycles are grouped by their order relative to the stationary-phase transition, and the division strategy is obtained from the relationship between birth size *s*_*b*_ and added size at division ⟨∆d⟩. For entry into stationary phase, cycles were indexed backward (from −6 to −1) from the last observed divisions before growth arrest. For exit from stationary phase, cycles were indexed forward (from 1 to 6) from the first complete divisions after growth resumed [Fig. 2(a)].

**FIG 2.**
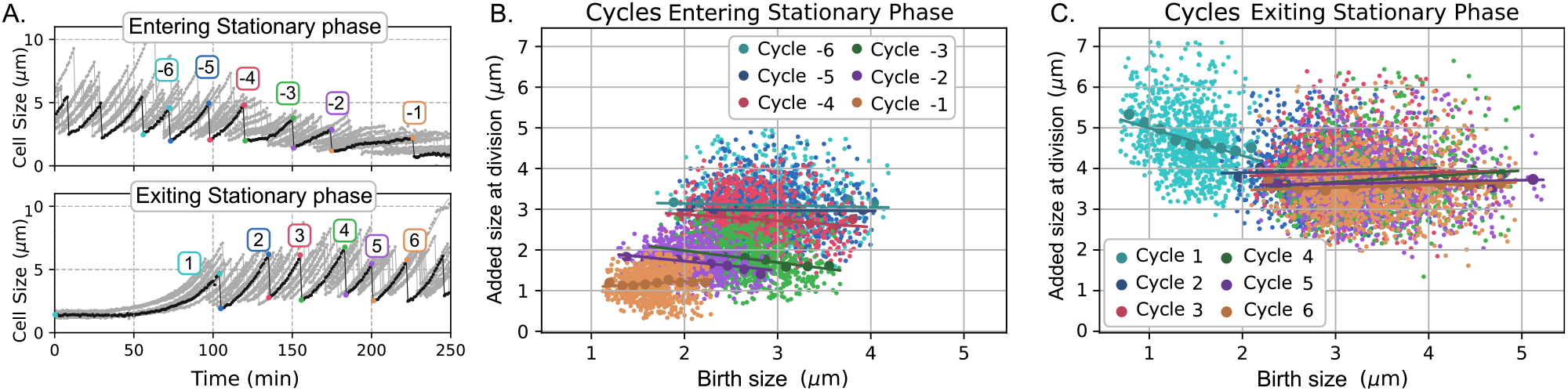
Division strategy across successive cell cycles during entry into and exit from stationary phase. (a) Cell size trajectories over time illustrating how complete cell cycles were indexed relative to the stationary-phase transition. Top: Cycles preceding entry into stationary phase were indexed backward from the last observed divisions before growth arrest. Bottom: Cycles following exit from stationary phase were indexed forward from the first complete divisions after growth resumed. (b) and (c) Division strategy during entry into and exit from stationary phase, respectively, illustrated by the relationship between added size (length) and birth size across successive cell cycles. A slope of zero indicates an adder strategy, while negative and positive slopes indicate sizer-adder and timer-adder strategies, respectively. In (b) and (c), colors indicate the cell-cycle groups defined in (a). Darker points and curves represent binned averages, while solid lines show linear fits indicating the dependence of added length on birth length.

During entry into stationary phase, the relationship

between added size and birth size changed across successive cycle groups [Fig. 2(b)]. Divisions before nutrient depletion (cycles −6 and −5) followed an adder strategy, with weak dependence of added length on birth length. Closer to stationary phase (cycles −4 to −2), the slope became negative, indicating a transient shift toward sizer-adder control, together with a progressive decrease in mean added size across cycles. In the final cycle before growth arrest (cycle −1), the mean added length was substantially reduced, while added length again showed almost no dependence on birth length, consistent with adder control.

Exit from stationary phase showed a different pattern [Fig. 2(c)]. The first complete cycle after growth resumed was characterized by small birth lengths and large added lengths, suggesting substantial size recovery before division. This first post-starvation cycle also showed a strong negative correlation between birth length and added length, consistent with sizer-adder control. In later cycles (cycles 2–6), the regression slope was close to zero (adder), while the mean added length gradually decreased from a relatively large value (≈ 4 *µ*m) toward the steady-growth value (≈ 3.5 *µ*m). Although this discrete cell-cycle analysis reveals trends across successive divisions, it averages division control over each complete cycle and therefore loses information about changes on shorter timescales. This limitation motivates the continuous-time inference framework, which can resolve changes in division propensity during rapid environmental transitions.

### B. Maximum-likelihood inference captures time-varying division parameters

We next applied the maximum-likelihood framework of Sec. II to infer, in each time interval, the mean added size at division⟨ ⟨∆d⟩ ⟩ , its noise 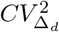 , and the strategy parameter *α* (*α* = 0 for an adder, *α <* 0 for a shift toward sizer-adder, and *α >* 0 for a shift toward timer-adder). Given the observed size trajectories and division events, we identified the parameter values that maximize the likelihood of the data, using the division probability of Eq. (6). We compared two models: an adder model, which assumes *α* = 0 (red line in Fig. 3), and the alternative model with *α* 0 (purple line in Fig. 3).

**FIG 3.**
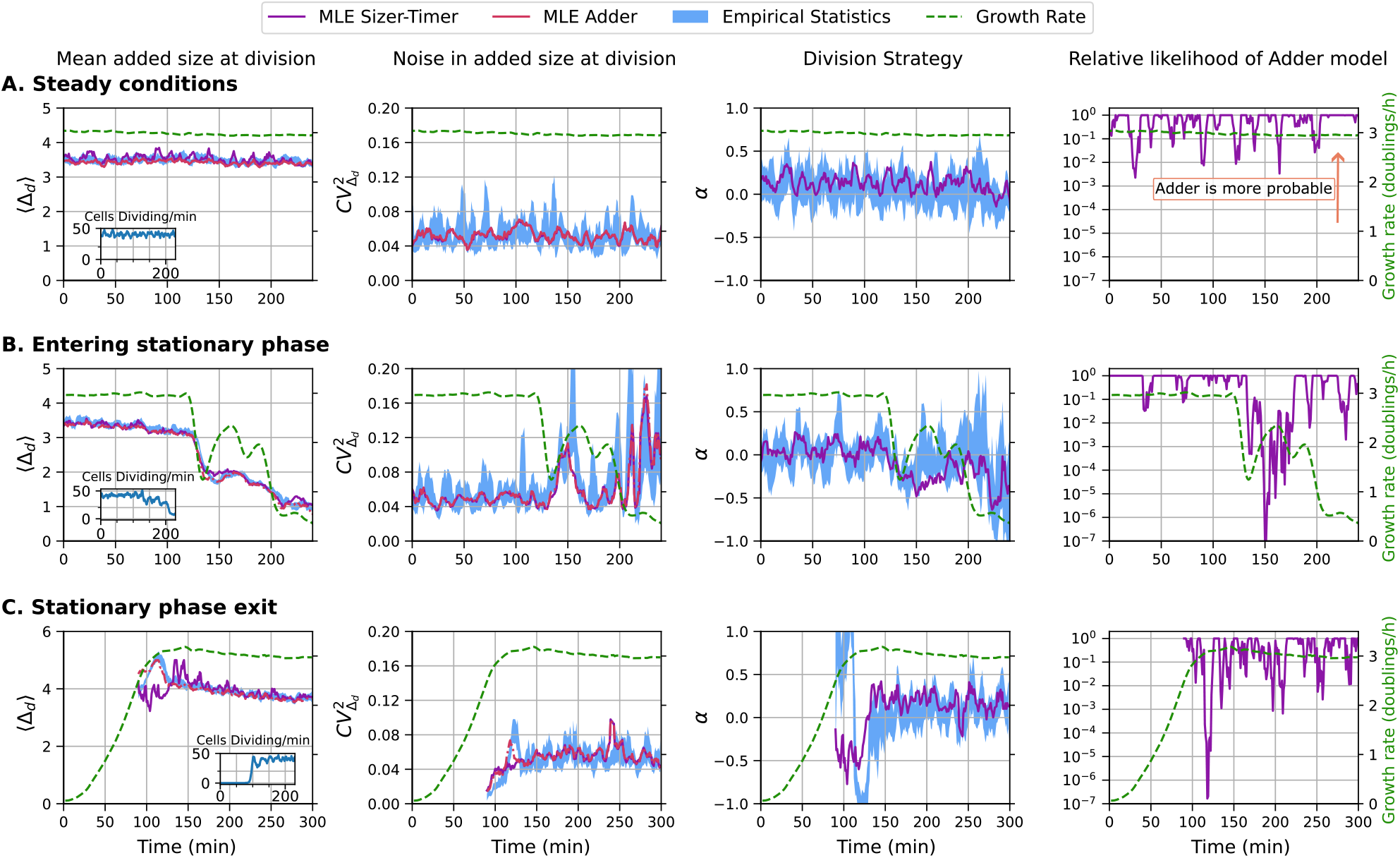
Dynamic inference of cell division parameters across different environmental conditions. Maximum-likelihood estimation [Eq. (8)] is used to infer division parameters from single-cell size trajectories over time. Rows correspond to three physiological conditions: (a) steady growth, (b) entry into stationary phase, and (c) exit from stationary phase. Across all rows, the four columns show, from left to right, the inferred parameters and model comparison over time. First column: Scale parameter, representing the mean added size at division ⟨ ∆_*d*_ ⟩ . Insets display the number of cells dividing per minute during the time-lapse. Second column: Shape parameter, describing noise in added size quantified by 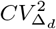. Third column: Division strategy parameter *α*, describing the linear relationship between birth size and added size in cell division control (adder: *α* = 0; sizer-adder: −1 *< α <* 0; timer-adder: 0 *< α <* 1). Fourth column: Relative likelihood of the adder model compared with the alternative division model, calculated using the Akaike information criterion [Eq. (12)]. Green dashed lines indicate the estimated mean growth rate, *µ*(*t*), in doublings per hour, with the corresponding axis shown on the right side of the fourth column. Red dash-dotted lines represent the parametric inference restricted to a pure adder strategy, while solid purple lines indicate the inference for the alternative model with an additional free parameter *α*. Blue shaded regions indicate the empirical parameter estimates (95% confidence interval) calculated directly from the subset of dividing cells in each time window by bootstrap resampling. Data from 952 single-cell lineages.

The alternative model has one more parameter; the trade-off between its better fit and its added complexity is quantified by the relative likelihood of the adder model [Eq. (12); fourth column in Fig. 3]. Values below 10^−3^ indicate substantially weaker support for the adder model relative to the alternative model.

Under steady growth conditions, the inferred parameters remained approximately constant over time for both the adder and the alternative models [Fig. 3(a)]. The inferred mean added size, ⟨ ⟨∆d⟩ ⟩ , and division noise, 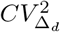 , were also similar for the two models. Notably, ⟨ ⟨∆d⟩ ⟩ follows a smoother trajectory under the adder model (*α* = 0) than under the alternative model. Although the inferred *α* for the alternative model was consistently positive, the relative likelihood of the adder model remained high, so the alternative strategy was not favored. As an empirical reference, we also computed the mean and noise of added size directly from cells dividing within each 3-min window (the current 1-min interval plus the preceding and following ones, to smooth the estimates). These model-independent values have relatively wide confidence intervals [blue regions in Fig. 3]. The maximum-likelihood estimates remained within these empirical confidence intervals, indicating that the inferred parameters are consistent with model-independent division statistics.

During entry into stationary phase, the estimated mean growth rate decreased over time as nutrients were depleted [green dashed line in Fig. 3(b)]. The decline in growth rate was not strictly monotonic but instead showed transient drops and recoveries. We associate these fluctuations with successive diauxic shifts, brief periods following the exhaustion of a preferred carbon source in the growth medium [55, 56]. During these shifts, cells temporarily slow or halt growth to adapt their metabolism to an alternative carbon source, a sequential process that continues until all available nutrients in the environment are depleted.

Additionally, during entry into stationary phase, the inferred mean added size decreased over time [Fig. 3(b)], accompanied by a reduction in the number of dividing cells [inset of Fig. 3(b)]. The noise in added size peaked repeatedly, mostly just after each dip in growth rate. The inferred strategy parameter *α* also shifted toward negative values during this interval, indicating a transient departure from pure adder division strategy toward sizer-adder control. In these periods, the relative likelihood of the adder model, *p* [Eq. (12)], decreased below 10^−3^, so the deviation from the adder is statistically supported. After the last strong decrease in growth rate, the relative likelihood of the adder increased again to values consistent with adder division. This pattern agrees with the discrete cell-cycle analysis shown in Fig. 2(b).

During exit from stationary phase, inference was limited by the long delay between fresh-medium addition and the first observed divisions. Cells grew for approximately 100 min before division events began [Fig. 3(c)]. Because the inference depends on observed division events, parameters were estimated only when at least five divisions occurred within a time interval. After growth resumed, the inferred mean added size was initially high, exceeding 4 *µ*m, and then decreased gradually over approximately 150 min (about six generations). The first post-starvation divisions showed lower noise than divisions during steady growth, suggesting more precise division control.

Comparison of the adder model (red) and the alternative model (purple) showed that the alternative model yielded a lower mean added size together with a negative strategy parameter, *α* ≈ −0.5, between 110 and 130 min. In this interval, corresponding approximately to the first post-starvation divisions, the relative likelihood of the adder model was very low (*<* 10^−6^), indicating strong support for a sizer-adder strategy. In subsequent divisions, the adder model was again favored, consistent with the discrete cell-cycle analysis in Fig. 2(c). The inferred *α* during the first division interval did not fully match the empirical estimate, likely because few divisions occurred in this window. This discrepancy suggests that strategy estimates are less reliable when the number of observed division events is small.

Together, the inferred parameters and model comparison show that the proposed framework can quantify how cell division control is modulated under time-varying growth conditions. Across all conditions, *E. coli* division remained broadly adder-like, but environmental transitions produced transient changes in the added size, division noise, and inferred division strategy. The strongest deviations occurred near entry into stationary phase and during the first divisions after exit from starvation, when cells shifted toward sizer-adder control. The maximum-likelihood estimates agree well with the empirical statistics of added size at division (mean and noise) and with its correlation with birth size (strategy). The approach thus provides a concrete way to quantify the scale and stochasticity of division in continuous time under nonsteady conditions.

## IV. DISCUSSION

In this study, we applied and extended a maximum-likelihood inference framework based on a coarse-grained model to quantify cell division control properties under time-varying growth conditions. By modeling division as a stochastic jump process with propensities dependent on cell size and added size, the framework infers three interpretable quantities: the effective mean added size at division ⟨∆d⟩, the division noise 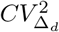 , and the division strategy parameter *α*. Unlike classical cell-cycle methods, which estimate division strategy from statistics over completed cycles, our approach is based on division events counted across a population of cells sharing the same environment. As a result, the division parameters are well defined even when the environment changes faster than a generation time.

The three inferred parameters map onto distinct aspects of cell-cycle physiology. The scale parameter ⟨ ⟨∆d⟩ ⟩ reflects the amount of biomass a cell commits to accumulating before septation, and is therefore a readout of the set point that couples resource allocation to the division machinery [57, 58]. The shape parameter 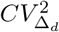 reflects the precision with which the cell size set point is enforced. Several mechanisms have been proposed to explain the variability in cell-size regulation. Under optimal growth conditions, division is coordinated with growth and size noise reaches a fixed level [49, 59]. A major source of randomness in division has been linked to the number of effective rate-limiting steps in the cell cycle [44, 51, 60]. An increase in this noise may therefore indicate changes in the synchronization of growth and division. Finally, *α* measures how strongly birth size is carried forward into the division decision, distinguishing initiation-limited control (sizer-adder, *α <* 0) from accumulation-limited control (adder, *α* = 0) [50, 61].

During entry into stationary phase, we observed a progressive decline in ⟨∆d⟩ that closely followed the decreasing growth rate. The initial drop in ⟨∆d⟩ coincided with the decrease in growth rate during the first diauxic shift. When the growth rate temporarily recovered, ⟨∆d⟩ followed with a modest increase, delayed by approximately 30 min. Subsequent decreases in ⟨∆d⟩ similarly corresponded to the growth rate drops of later diauxic shifts. Mechanistically, this decline is likely driven by differences in the production of division proteins (such as FtsZ and its regulators) related to the cell’s global translational capacity [57, 62, 63]. Because the proteins required for division accumulate faster than the cell can double its biomass, cells divide at a progressively smaller size. The increase in 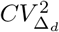 at each drop in growth rate is consistent with the idea that nutrient depletion changes the synchronization between biomass accumulation and division. Diauxic shifts require the cell to rewire carbon metabolism, and during this rewiring, flux into the division pathway varies considerably from cell to cell [64, 65]. The transient shift toward negative *α* during these windows also admits a simple interpretation: when biosynthesis slows abruptly, cells that were born large are closer to any absolute size or DNA-content threshold and divide sooner, whereas cells born small must accumulate disproportionately more. Under starvation, division becomes limited by reaching a critical state (e.g., completion of replication or a threshold divisome concentration) rather than by accumulating a fixed increment [66]. Sizer-adder control under slow growth has been reported in several systems, and our framework shows that it can arise transiently within a single nutrient downshift rather than only as a property of distinct steady states [67, 68].

Exit from stationary phase produced the largest deviation from adder division. Cells grew for roughly 100 min before the first division and then divided after adding an unusually large increment (*>* 4 *µ*m), with negative *α* (≈ −0.5). This is consistent with a recovery process in which biomass accumulation and division are temporally uncoupled: starved cells may first need to restore ribosome production, membrane and cell-envelope synthesis, and DNA replication before assembling the divisome and resuming division [62, 69]. Division in this window is then likely gated by the attainment of a competent physiological state rather than by an incremental size counter, which is consistent with a negative *α* [70].

The observation that the first post-starvation divisions have relatively low noise 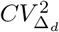 is notable, since noise usually rises during transitions. One interpretation is that when division is gated by a threshold that all cells must cross (for instance, replication initiation), the threshold itself acts as a noise filter: initial size heterogeneity accumulated during starvation is compressed rather than propagated, because small cells simply grow longer before reaching the commitment size and then add a relatively constant size before dividing [50, 71, 72]. The subsequent slow relaxation of ⟨∆d⟩ over roughly six generations then represents the gradual re-establishment of steady-state proportionality between growth and division.

These results have two implications. First, the adder may be best understood as the steady-state limit of a more general control law rather than as a fixed mechanism. Cells need not measure added size *per se*; rather, adder division strategy emerges when the accumulation of division-limiting components is proportional to growth rate [33, 59]. When that proportionality is disrupted, as during entry into stationary phase, the effective strategy deviates from the adder in the direction predicted by the limiting process. Second, transient sizer-adder control can affect cell proliferation. At entry into stationary phase, sizer-adder control makes large cells divide sooner, which increases cell number and reduces size variability within the clonal population [73]. We observe sizer-adder division mainly when cells recover from growth slowdowns, either during resuscitation from starvation or after a diauxic shift; in both situations, faster division of large cells could favor scavenging of newly available nutrients and survival.

Our model is phenomenological and identifies effective, not molecular, parameters. The specific form of the propensity function was chosen for numerical convenience [46, 48], and although the inferred moments are robust to this choice under steady growth conditions, we cannot exclude the possibility that some of the inferred *α* dynamics reflect model misspecification during rapid transitions. The inference also treats cells within a time window as independent and as sharing common parameters, which neglects lineage correlations and any persistent cell-to-cell heterogeneity in cell division control [74]. Finally, the framework requires division events within each window, so its temporal resolution is limited by the frequency of division events in the population. Accurate parameter estimation also requires a relatively large number of division events: in our analysis, each time window (1 min)contained approximately 50 divisions among the 952 tracked lineages. In our benchmark, we found how the error of the inference decays with the square root of the number of division events [46].

The formalism can be extended to asymmetric division and nonexponential growth by substituting *µs* for *ds/dt* in Eq. (2). [33]. Such extensions would make it applicable to organisms such as yeast and mammalian cells, where growth and division mechanisms differ from those of *E. coli* [31, 75, 76]. Our method is also robust to other sources of variability, such as noise in partitioning or growth rate, which strongly bias methods that rely solely on cell-size distributions rather than on division events [44, 47, 77]. Extending the propensity to depend on additional variables such as cell width, gene expression, or cell-cycle reporters is a straightforward generalization of the likelihood and would allow formal tests of the relevance of other variables [43]. Finally, embedding the inferred time-dependent division rules into population-level models would connect single-cell control to colony growth and lag-phase distributions, linking the parameters measured here to ecologically relevant fitness outcomes [28, 78–80].

## ACKNOWLEDGMENTS

We acknowledge support from NIH-NIGMS through Grant No. R35GM148351. We thank Somenath Bakshi (University of Cambridge) for contributing the dataset used in this study.

## DATA AVAILABILITY

The data set and data-processing scripts are publicly available on Zenodo. https://doi.org/10.5281/zenodo.22268346 [81].

## AUTHOR CONTRIBUTIONS

- **Conceptualization:** C.N., S.R.
- **Investigation:** S.R., C.N.
- **Data Curation:** C.N.
- **Formal Analysis:** S.R., C.N.
- **Funding Acquisition:** A.S.
- **Methodology:** S.R., C.N.
- **Resources:** A.S.
- **Software:** C.N.
- **Supervision:** A.S., C.V.
- **Visualization:** C.N., S.R.
- **Writing – Original Draft:** C.N., S.R.
- **Writing – Review & Editing:** C.V., A.S.

